# Concurrent separation of phase-locked and non-phase-locked activity

**DOI:** 10.1101/2022.03.14.484222

**Authors:** Shubham Singhal, Priyanka Ghosh, Neeraj Kumar, Arpan Banerjee

## Abstract

Brain dynamics recorded via electroencephalography (EEG) is conceptualized as a sum of two components, “phase-locked” and “non-phase-locked” to the stimulus. Both activities are understood to be stemming from different neuronal mechanisms and hence accurately characterizing them is of immense importance in neuroscientific studies. Here, we discuss the drawbacks of currently used methods to separate the phase-locked and non-phase-locked activity and propose a new method that decomposes the two components simultaneously. First, we demonstrate that single-trial separation of phase-locked and non-phase-locked power is an ill-posed problem. Second, using simulations where ground truth validation is possible, we elucidate the drawbacks of the widely used averaging method and efficacy of the proposed concurrent phaser method (CPM). Using two experimental datasets, audio oddball EEG data and auditory steady-state responses (ASSR) we show how the empirical signal-to-noise estimates warrant the usage of CPM to separate phase-locked and non-phase-locked activity.

## 1 Introduction

Previous studies modeled electroencephalogram (EEG) activity as a superposition of two independent components, one evoked and the other induced activity [Pfurtscheller and Da Silva, 1999]. The evoked activity was shown to be phase-locked to the stimulus, whereas induced activity was non-phase locked [Tallon-Baudry et al., 1997]. Studies have shown some stimuli only to evoke phase-locked activity, some to only induce non-phase-locked activity [Tallon-Baudry et al., 1997, Tallon-Baudry and Bertrand, 1999, Pfurtscheller and Da Silva, 1999]. In contrast, some categories of stimuli have been argued to elicit phase-locked (PL) and non-phase-locked (NPL) activity simultaneously [Deiber et al., 2007] [Zhang et al., 2020]. Typically, PL activity is analyzed in the time domain, whereas power in the frequency domain is used to measure NPL activity. Pfurtscheller and colleagues [Pfurtscheller and Da Silva, 1999] considered PL activity - transient response of pyramidal neurons triggered by a specific stimulus, whereas NPL activity to result from an increase or decrease in synchrony of ongoing background brain activity [Pfurtscheller and Da Silva, 1999]. The stimulus can change one or more parameters that control the oscillations in neuronal structures, which can change the strength of synchrony or desynchrony of the background activity [Pfurtscheller and Da Silva, 1999]. From a measurement perspective, synchrony or desynchrony are reflected in changes in the spectral power in one or more frequencies. Thus, any stimulus can contribute to an alteration of both PL and NPL activity simultaneously. Phase-locked and non-phase-locked activity represents distinct underlying cognitive and neural processes [Gruber et al., 2006].

Jervis and colleagues [Jervis et al., 1983] devised a statistical test (Rayleigh test) to evaluate the strength of phase clustering across trials and separate the EEG signal corresponding to a stimulus into two classes, one phase ordered and the other with non-significant phase ordering. The phase-ordered signal is captured by taking the average across trials, and defined as the event-related potential (ERP). In contrast, non-phase-locked activity is analyzed in the frequency domain by taking Fourier transforms. Kalcher and Pfurtschellar [Kalcher and Pfurtscheller, 1995] were the first to consider stimulus to generate both activities simultaneously and proposed the inter-trial Variance method (IVM) to separate phase-locked from non-phase-locked activity. In IVM, ERP power is subtracted from each trial power and an average of the squared residuals is interpreted as non-phase-locked activity. The non-phase-locked component at 40Hz has been shown to play a crucial role in visual feature binding [Tallon-Baudry and Bertrand, 1999]. IVM method works under the assumption that ERP is amplitude and shape invariant. Most studies do not verify this assumption. ERP varies trial-by-trial due to task practice, fatigue, attentional lapses [Kappenman and Luck, 2011]. It is well-known that ERP amplitudes decreased while increasing trial numbers [Knuth et al., 2006]. Some reports beforehand characterize the signal to be either made of predominantly phase-locked or non-phase-locked activity using phase-locking factor and Rayleigh test [Jervis et al., 1983]. However, to date, no method has been proposed that considers phase-locked and non-phase-locked activity co-occurring and varying with each trial, i.e., under the general condition-ERP amplitude varies across trials [Cohen and Donner, 2013].

In this study, we propose a method, which we refer to as the concurrent phaser method (CPM) that estimates both PL and NPL concurrently with the consideration of both varying trial-by-trial. Through simulations, we identify the limit case scenarios where the proposed CPM approach is equivalent to using the currently used averaging method. We also demonstrate where decomposition into PL and NPL at a trial-by-trial becomes ill-posed; rather only computation of the distributions of PL and NPL is possible. Most importantly, all the previously used methods analyzed one activity at a time, i.e., either PL or NPL. Both activities are considered co-occurring in CPM, and estimators are calculated concurrently. Finally, the dependence of averaging method PL and NPL estimates on the ratio variance of PL and NPL was demonstrated using two different EEG datasets.

## 2 Methods

### 2.1 Averaging method to compute PL and NPL components

The prevalent method of separating phase-locked from non-phase-locked power is the averaging method [Truccolo et al., 2002] [Siegel and Donner, 2010], [Klimesch et al., 1998], [Donner and Siegel, 2011], [Cohen, 2014]. The event-related potential(ERP) is calculated by averaging time series across all trials. This ERP is subtracted from each trial time series and converted to frequency domain(via short-term-Fourier-transform or using Wavelet transform); average across all trials in the frequency domain is considered as time-frequency representation of non-phase locked activity. The phase-locked power for each trial is calculated by subtracting non-phase locked power from the total power of the signal. The pre-stimulus or post-stimulus activity is considered the baseline to which both activities are statistically tested. In **Figure(1)**, panel (A) shows the steps to get NPL and PL power using the averaging method. Where, *S*_*r*_(*t*) is a single-trial signal, ⟨. ⟩ represents the average across trials, TF is the Time-Frequency transform, subscript ‘r’ represents *r*^*th*^ trial activity. We simulated trials of 2 sec at 0.001-sec resolution at 10Hz. The signal has an only phase-locked component for 0-0.5sec, an only non-phase-locked component for 0.5-1 sec, simultaneous PL and NPL components for 1-1.5sec and for 1.5-2sec both PL, NPL powers are increased four-folds. We can see in **Figure (a3) of (1)** that although there is no NPL power from the 0-0.5sec but averaging method is showing some pawer as evident from the plot **(a5)**. Panel **(C)** of **Figure(1)** shows the algorithmic steps to CPM proposed in this study. Comparing **(a5)** and **(c4)**, we see that CPM does not show any NPL power, whereas the averaging method shows NPL power for 0-0.5sec. Comparing two for times 1-2sec, we can see that NPL power is overestimated in averaging method. The reason for these discrepancies is discussed in section 2.1.

**Figure 1:**
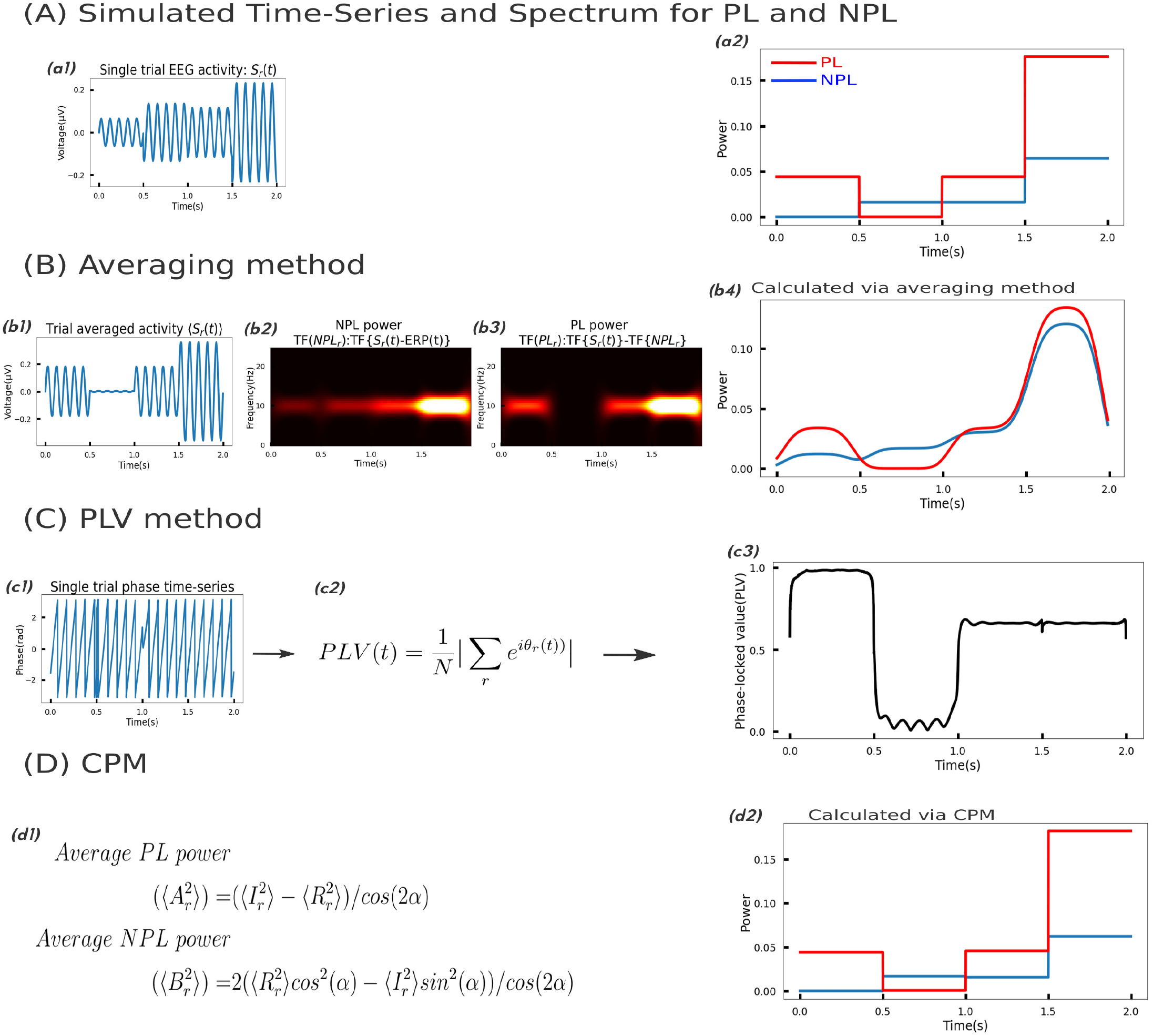
Signal simulated at 10Hz with only phase-locked activity for 0-500ms, only non-phase-locked activity for 500-1000ms, simultaneous PL and NPL for 1000-1500ms and for 1500-2000ms, both PL and NPL power are quadrupled. The simulated signal is analyzed with the averaging method, phase-locking value(PLV), and CPM. (A) Simulated single-trial time series and power for each time point. (B) Algorithmic steps to Averaging method, (b1) signal averaged across all trials to get ERP, (b2) NPL power, time-frequency representation of (*S*_*r*_(*t*)−*Erp*), (b3) PL power, (b4) PL, NPL power estimate varying with time at 10Hz. (C) Algorithmic steps for PLV method, (c1) phase time-series is calculated for each trial(calculated using analytical signal), (c2) phase locked value defination, (c3) PLV varying with time is calculated. (D) Algorithmic steps to CPM, time-series into n bins(n is time steps of *A*_*r*_, *B*_*r*_ calculation, here (n=4), (d1) Average PL, NPL power calculated from Fourier coefficient at 10 Hz, (d2) PL, NPL power calculated using CPM estimates.

There are variates of this method, but the central idea is averaging out the non-phase-locked activity and assuming phase-locked activity to be trial invariant, which remains the same across all the variant methods, reported to date. IVM intertrial variance method is a time-domain analysis, where time series is first bandpass-filtered according to the frequency of interest and then averaged across all trials. Each bandpass filtered time series is subtracted with the calculated average and squared to get the power-time evolution[Kalcher and Pfurtscheller, 1995]. BW. Jervis et al. proposed a method to measure if the signal is predominantly phase-locked or non-phase-locked, proposed phase ordering factor and Rayleigh test[Jervis et al., 1983]. The paper accepts that it is impossible to establish which explanation applies (additive or phase-reordering) based on proposed methods when the additive energy is small[Jervis et al., 1983]. All the studies of separating phase-locked and non-phase-locked activity are done under the assumption that ERP is trial invariant[Tallon-Baudry and Bertrand, 1999][Cohen and Donner, 2013][Deiber et al., 2007][Zhang and Han, 2021][Fu et al., 2011]. We relax the trial-invariant PL activity assumption(at an interested frequency) and calculate PL, NPL amplitude concurrently. We also analyze the limiting case where the above method is valid.

### 2.2 Phase Locked Value (PLV)

In this section, we discuss phase locked value as a method used to quantify the phase-distribution across trials. A uniform phase distribution from 0 to 2*π* gives a PLV of zero, whereas a single phase across all trials gives a PLV of one. Panel**(B)** of **Figure 1** shows the steps involved in PLV analysis, at first signal is filtered at the frequency of interest, then phase-time series is calculated (using analytic signal) for each trial. Phase locked value is calculated at each time point, defined as [Lachaux et al., 1999]

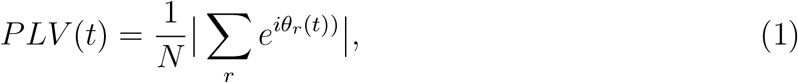

where N is the total number of trials, ‘r’ represents *r*^*th*^ trial, *θ*_*r*_(*t*) is the *r*^*th*^ phase at time t. PLV varies from zero to one for no synchrony to complete synchrony. PLV method can be viewed as an indirect method of calculating phase-locked power, though it does not calculate PL or NPL power. As shown in **(b3)** of **Figure 1**, PLV is 1 for 0-0.5sec where there is only phase-locked activity and zero for 0.5-1sec for which there is only non-phase-locked activity. We have a PLV of 0.6 for 1-2 sec, which does not change as PL and NPL power are increased for 1.5-2 sec. The reason for no change in PLV value even though PL and NPL power are changed is discussed in section 3.1. PLV does not calculate NPL or PL power, but it depends on NPL to PL content in the signal. As shown in **Figure 2**, phase locked value decreases as NPL content increases with respect to PL content. We get a PLV of 1 for signal with no phase-locked content i.e., pure phase-locked signal, and PLV decreases to zero as the NPL to PL power ratio increases. The phase locked value is used to make the phase-synchorony comparison between rest and stimulus-response, and an indirect inference for PL and NPL power is made through PLV as in [Goffaux et al., 2004].

**Figure 2:**
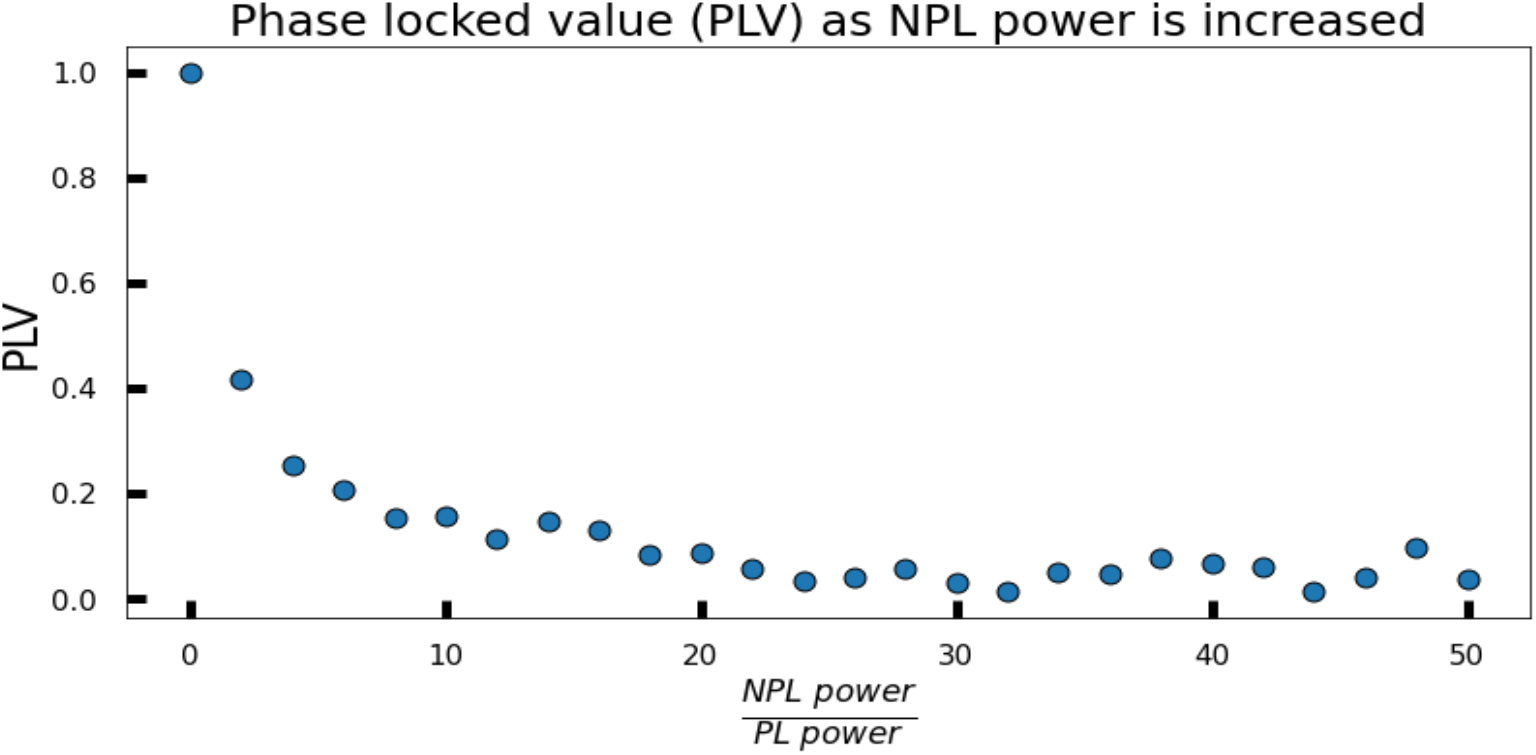
Phase locked value for varying NPL to PL content in the signal.

### 2.3 Concurrent phaser method (CPM)

Consider a signal snippet from 0 to *T* time. The signal is represented as the sum of sine functions with frequencies(0, 1*/T*, 2*/T*, 3*/T…*.)

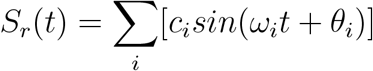

where *ω*_*i*_ ∈ [0, 2*π/T*, 4*π/T*, …]. We assume the signal to have phase-locked and non-phase-locked activity at each frequency. Both PL and NPL components are considered time-locked. We assume the phase of PL activity to be trial-invariant and NPL activity to vary across trials. Therefore, the signal can be written as equation(2)

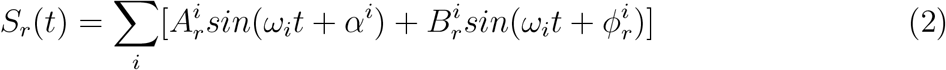

Where 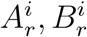 are trial varying non-negative phase-locked and non-phase-locked amplitudes simultaneously, *α*^*i*^ is the phase of the phase-locked part(constant across trials), 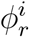 is the trial-varying phase of non-phase-locked part at frequency *ω*_*i*_ with the signal decomposed into frequencies *ω*_*i*_’s. Now, considering signal at a certain frequency *ω*_*o*_, equation(2) can be written as equation(3).

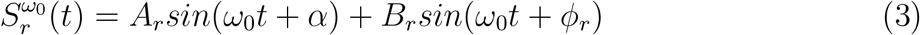

From here on, we will only consider signal at a specific frequency, and 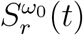 will be referred to as *S*_*r*_(*t*). 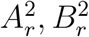 are the PL, NPL power at *r*^*th*^ trial. We model signal as given by equation(3) and try to calculate *A*_*r*_, *B*_*r*_ in section 3.2 and 3.3.

### 2.4 Generation of simulated data

Throughout the study, data is simulated to test the averaging method estimates, verify the CPM estimators and show the difference between averaging and CPM. The data was simulated according to equation(3), for all estimation purposes. We simulate the signal of 1 second at a resolution of 1ms at 10 Hz. We simulate *A*_*r*_ and *B*_*r*_ to have a normal distribution with a specified mean and standard deviation. The trial-varying phase *ϕ*_*r*_ was taken from a uniform distribution (U[−*π, π*]), *α* is chosen or varied between -*π* to *π*. A thousand trials were simulated for each specific value of *A*_*r*_ or *B*_*r*_ means, standard deviation.

## 3 Results

### 3.1 Drawbacks of averaging method and PLV method

#### 3.1.1 Averaging method

We assume the signal to have only one frequency i.e., *ω*_0_ with PL phase constant and NPL phase varyies across trials as in equation(3). We calculate PL and NPL powers following the procedure described in section 2. (′⟨. ⟩ ′ represents trial average).

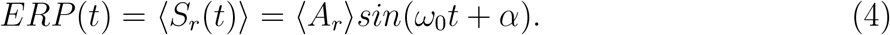

Non-phase-locked Power(with power given by square of fourier coefficient):

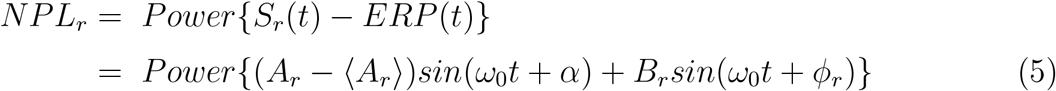

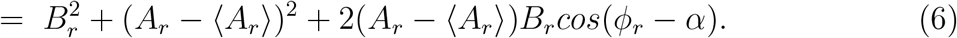

From equation 3 we can compute the total power

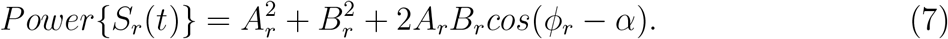

Thereby, the PL power (*PL*_*r*_) can be expressed as subtraction of NPL power from the total power.

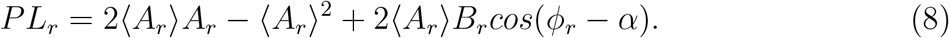

As shown in equation (6) and (8), *PL*_*r*_ or *NPL*_*r*_, are composite functions of *A*_*r*_ and *B*_*r*_ and are inseparable. Hence, theoretically, *PL*_*r*_ and *NPL*_*r*_ cannot be seperated. Therefore the widely used averaging method needs to be revisited for the general case of trial-varying phase-locked activity.

We simulate signal considering equation (3) for 1000 trials and estimate PL, NPL powers via the averaging method, **Figure(3)** shows the results. The figure clearly shows that the averaging method fails to capture the single-trial phase-locked and non-phase-locked power. Neither the PL nor NPL power falls on a linear line. Some of the PL power estimates are negative, clearly showing the loop-hole in the procedure. We calculated a low Pearson correlation of 0.27 between estimated and simulated NPL power(**Figure(3)[B]**). This makes NPL power estimate at single-trial level unreliable to correlate with any trial-level parameter. However, NPL power estimate at the single-trial level is correlated with reaction times by [Cohen and Donner, 2013],[McKewen et al., 2020]. The reported correlations have low value and should be scrutinized carefully before drawing any conclusion, as single-trial averaging method estimates are not accurate.

**Figure 3:**
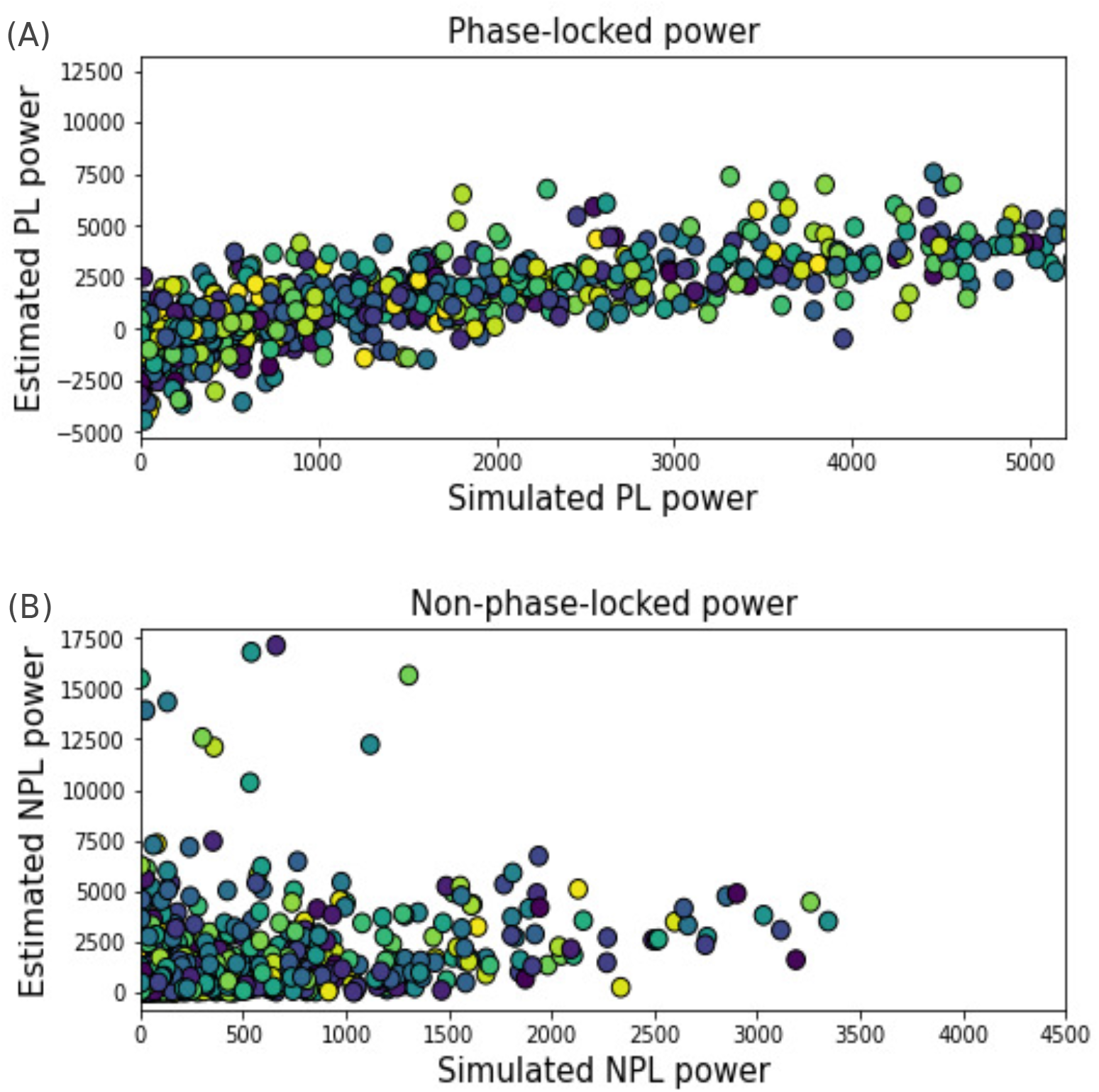
Single-trial PL, NPL power calculation via the averaging method. (A) phase-locked power, simulated vs estimated, (B) non-phase-locked power, simulated vs estimated (Pearson correlation(r)=0.27).

It is evident from equations(6) and (8) that variation in PL power corrupts the NPL and vice-versa, which should not be the case for the ideal separation of PL and NPL. As can be seen in the **(a5)** of **Figure 1** for time 0-0.5 sec where there is no NPL power, variance in PL activity is estimated as NPL power. And even for time 1-2 sec variance in PL activity corrupts NPL power estimates.

#### 3.1.2 PLV method

It is observed that phase locked value calculation is independent of the variance in PL or NPL activity. PLV method has a limitation that it does not calculate PL or NPL power, but it depends on ratio of NPL to PL power. This renders the PLV method ineffective for simultaneous change in both PL and NPL power cases. As can be seen in **(b3)** of **Figure 1**, though PL and NPL power have increased for 1.5-2 sec relative to 1-1.5 sec but PLV remains unchanged. PLV method will give an erroneous conclusion in case of simultaneous change in PL and NPL power, even if the increase is not proportional. PLV measures one content w.r.t. to the other; if both changes from one task to the other, the comparison between tasks is erroneous.

### 3.2 Single-trial separation of phase-locked and non-phase-locked power: an ill-posed problem

We model the signal as the form given in equation(3). The goal here is to calculate *A*_*r*_,*B*_*r*_,*ϕ*_*r*_. The Fourier coefficient of the signal at frequency *ω*_0_ is given by

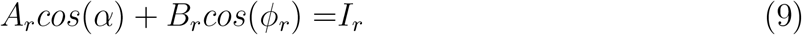

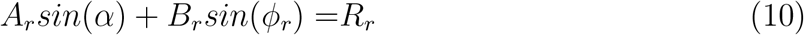

where *I*_*r*_ and *R*_*r*_ are the real and imaginary parts of Fourier coefficient. *α* is a constant phase (phase-locked phase) at frequency *ω*_0_, the estimate of *α* is given by *tan*^−1^(⟨*R⟩ / ⟨I⟩*) (for calculation, see in Appendix: A). Equations (9) and (10) form an undetermined non-linear equations system. We have three unknowns(*A*_*r*_, *B*_*r*_, *ϕ*_*r*_) and two equations. Therefore, there ia an infinite number of solutions for *A*_*r*_, *B*_*r*_, *ϕ*_*r*_ for a given value of *R*_*r*_ and *I*_*r*_. The equations (9) and (10) can be reduced to

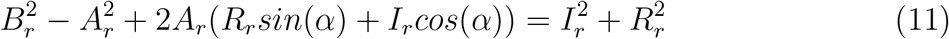

For a given value of *R*_*r*_,*I*_*r*_,*α*, the solution space of *A*_*r*_, *B*_*r*_ can be seen graphically, shown in **Figure(4)**, which shown infinite solutions for *A*_*r*_ and *B*_*r*_.

**Figure 4:**
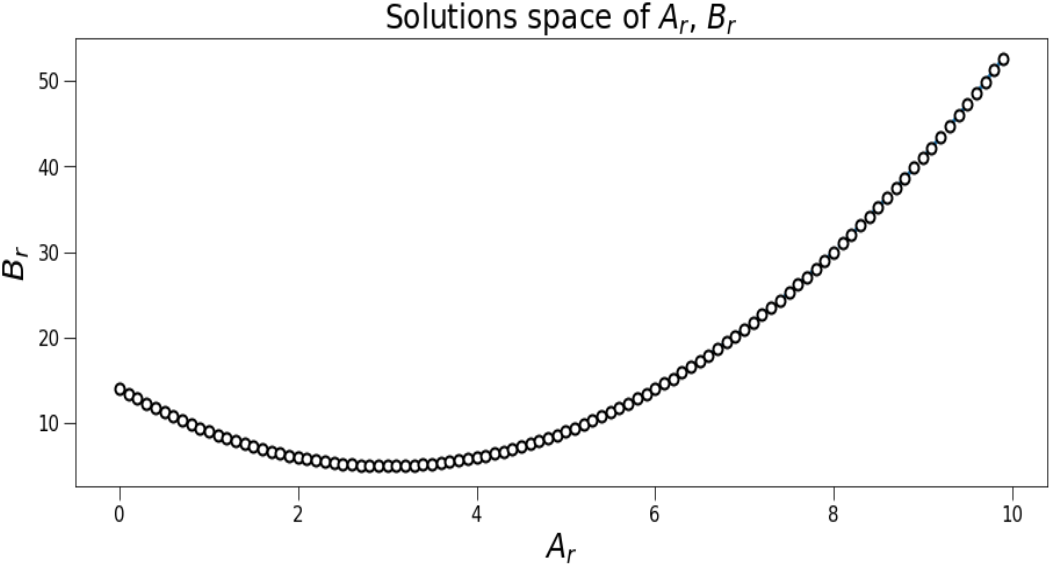
Solution space of *A*_*r*_, *B*_*r*_ for *R*_*r*_=2.24, *I*_*r*_=3, *α*=0.

### 3.3 Mean and Variance of PL and NPL power at group level using CPM

We observed the indeterministic nature of separating phase-locked from non-phase-locked power at the single level in the preceding section. We probe the question, can phase-locked and non-phase-locked activity be separated at the group level, i.e., combining all the trials. We set to find the mean and variances of phase-locked and non-phase-locked amplitudes to statistically comment about the separation of PL and NPL activity within and between groups of trials.

Taking averages of equations (9), (10) and considering *ϕ*_*r*_ to be uniformly distributed gives

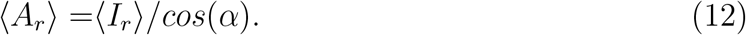

Substituting *A*_*r*_ from equation (9) into equation(10) gives

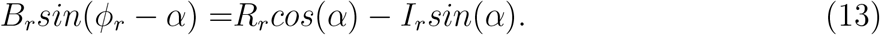

Since *B*_*r*_ is non-negative and sin(*θ*) has a positive value for *θ* ∈ (0, *π*), we average equation(13) for trials where the right-hand side, i.e., *R*_*r*_*cos*(*α*) − *I*_*r*_*sin*(*α*) = *Z*_*r*_ is positive. For trials where *Z*_*r*_ is positive, ⟨*sin*(*ϕ*_*r*_ − *α*) ⟩ = 2*/π*, which gives

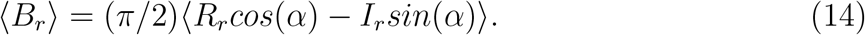

Equations (12) and equation (14) give the estimate of mean phase-locked and non-phase-locked amplitude. For calculating the variances, we square equations (9) and (10) to get

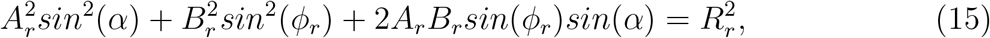

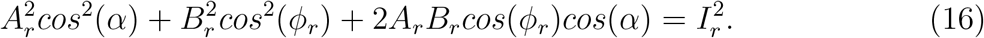

Averaging and adding equations (15), (16) gives

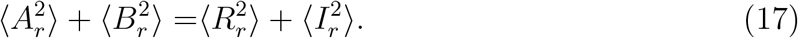

Averaging equation (16) gives

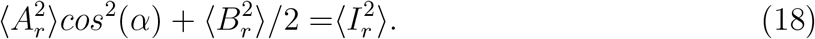

Solving equations (17),(18) for 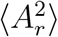 and 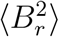, we have

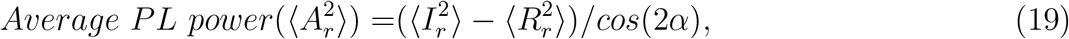

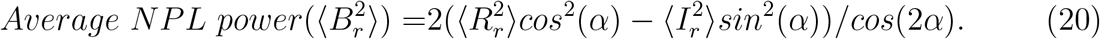

In conclusion, means and variances of *A*_*r*_, *B*_*r*_ are given by

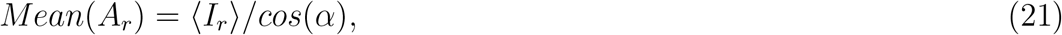

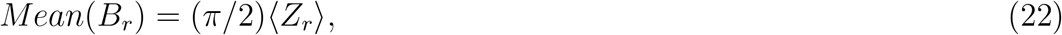

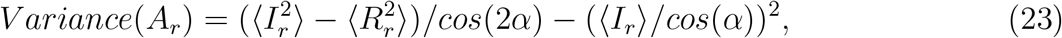

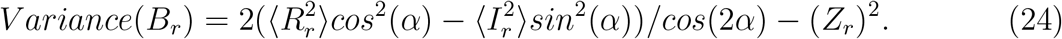

where *Z*_*r*_ = *R*_*r*_*cos*(*α*) − *I*_*r*_*sin*(*α*) for trials with *Z*_*r*_ positive.

These estimates of means and variances are used to make statistical inferences to separate phase-locked from non-phase-locked activity within and between tasks.

### 3.4 Estimator verification through simulation

In this section, we verify the estimators proposed in the previous section through simulations. We simulated 1000 samples of varying phase-locked, non-phase-locked amplitude, and non-phase-locked phase(*ϕ*_*r*_) for each estimate. We varied the simulated mean and variances of phase-locked, non-phase-locked amplitude and tracked their estimates. To see the robustness, we varied one parameter (e.g. *A*_*r*_ mean) while keeping all other parameters fixed (i.e., *A*_*r*_ variance, *B*_*r*_ mean, *B*_*r*_ variance). **Figure 5** shows the estimated versus simulated mean and variances. The top panels (a,b) in **Figure 5** which represents the estimates of mean, show that estimates lie on the straight line with negligible error, i.e., proposed estimators are good calculations of the simulated means. The lower (c,d) representing the estimates of variances shows that estimates follow the linear line but have a larger error than the mean estimates. We conclude that both mean and variance estimators are reasonale estimates.

**Figure 5:**
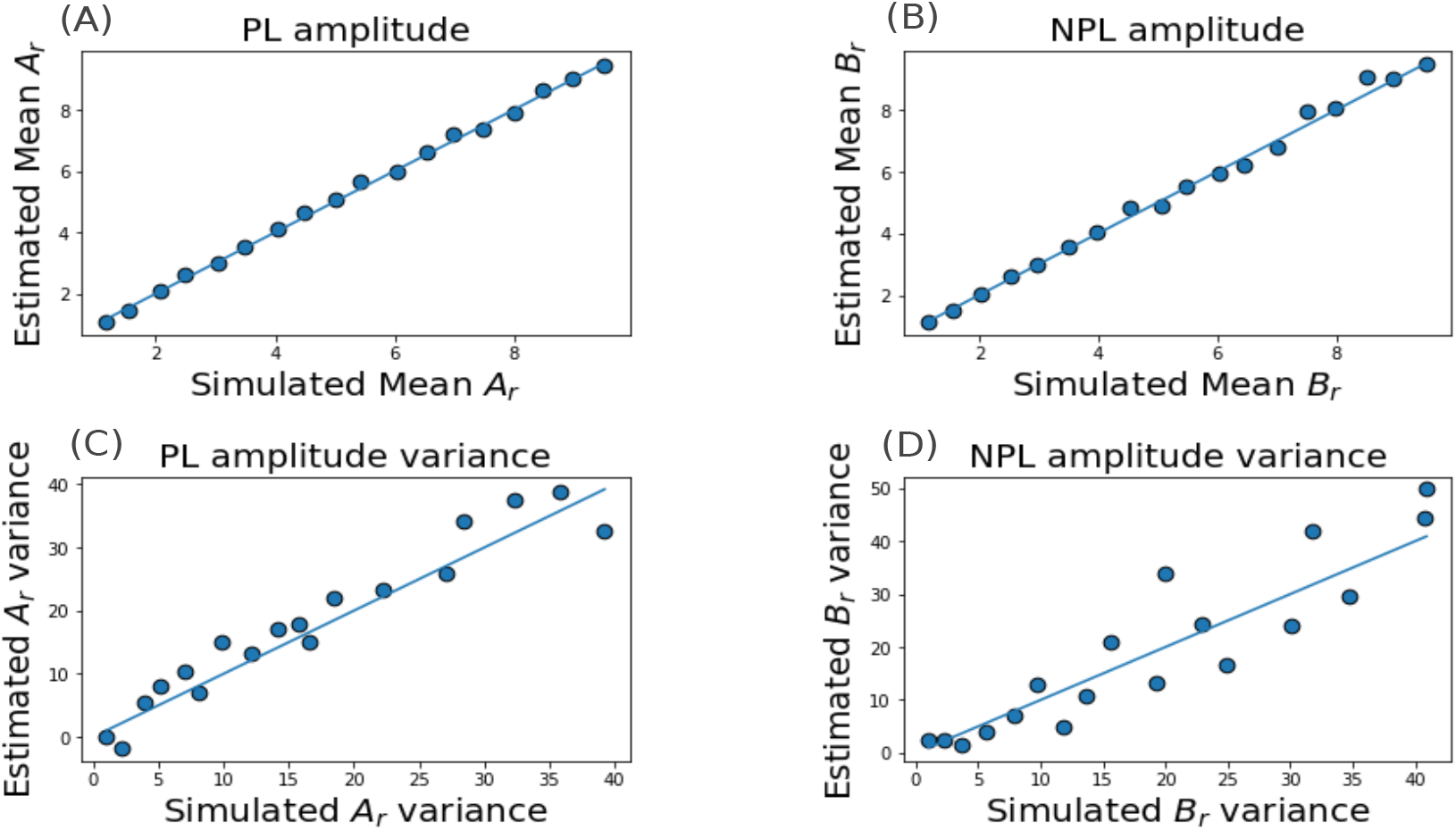
Verification of the proposed CPM estimators, estimated (A) *A*_*r*_, (B) *B*_*r*_ versus simulated mean, estimated (C) *A*_*r*_, (D) *B*_*r*_ variance versus simulated *A*_*r*_, *B*_*r*_ variance.

### 3.5 Application on Simulated Data

In this section, we first compare the estimates of CPM versus that of the averaging method and show that variation in phase-locked activity leaks into the estimate of non-phase-locked activity in the averaging method. In contrast, it does not affect the non-phase-locked activity estimates in CPM. Second, we show that CPM correctly identifies the statistical difference between the distributions of phase-locked or non-phase-locked activity between tasks, whereas averaging method gives erroneous results. The NPL power at single-trial calculated via averaging method is given by equation (5), taking averages on both sides, average NPL power is

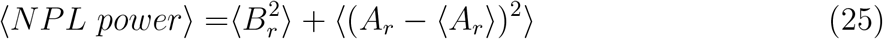

Non-phase-locked power estimate using averaging method shows dependence on the phase-locked activity, i.e., the variation in phase-locked statistics can leak into the estimate of non-phase locked activity leading to erroneous conclusions.

It can seen from equation(25) that large *A*_*r*_ variance i.e., ⟨ (*A*_*r*_ − ⟨*A*_*r*_ ⟩)^2^⟩ w.r.t 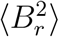 will affect NPL power estimate (via averaging method). So we simulate trials with varing *A*_*r*_ variance(keeping *B*_*r*_ means and variances constant) and estimate NPL power via CPM and averaging method. **Figure 6** shows non-phase-locked power estimates varies with the varying *A*_*r*_ variance(as 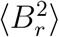 is constant). The CPM estimator of NPL power is not affected by the increase in the variance of *A*_*r*_. Averaging method estimate of NPL power increases linearly with an increase in *A*_*r*_ variance, i.e., the variance in *A*_*r*_ corrupts the NPL power estimate. So, we conclude that averaging method does not correctly separate the phase-locked power from non-phase locked power, as variation in one affects the other, even at the trial-average level. It is important to note here that CPM approaches averaging method estimate for low phase-locked amplitude variance. For 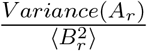 less than 0.5 CPM and averaging method estimate of average NPL power are approximately equal.

**Figure 6:**
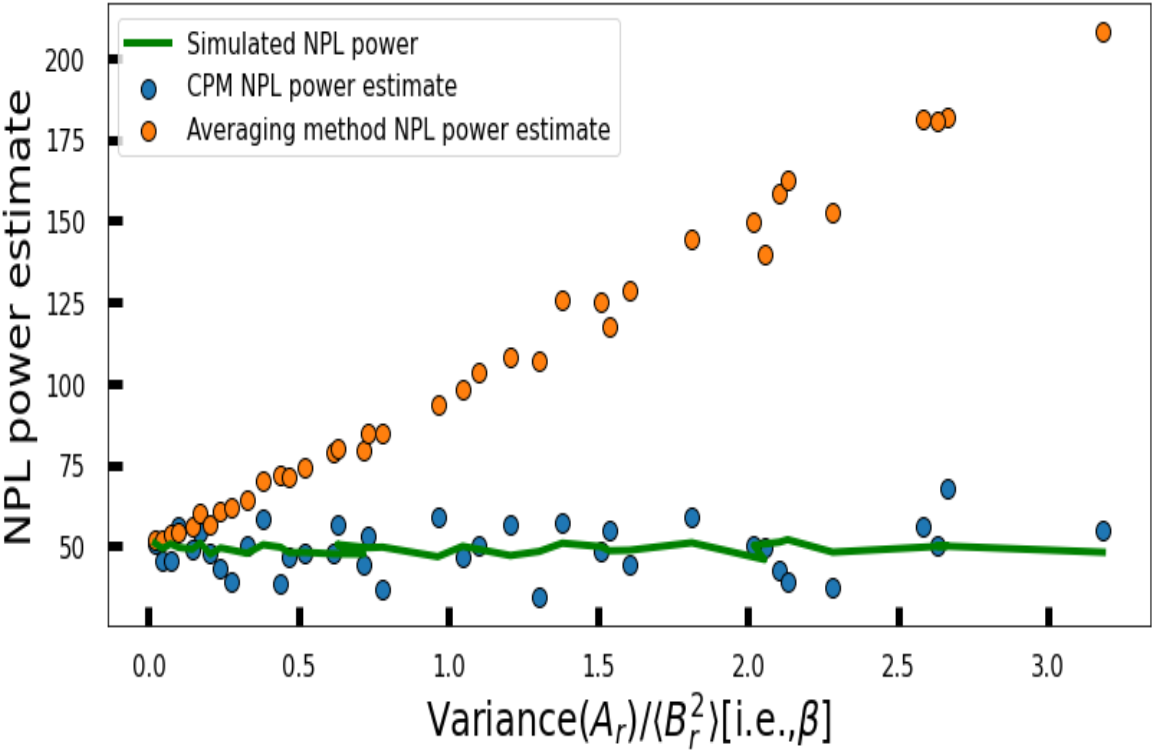
Estimated NPL power, averaging method(orange), CPM estimate(Blue) and simulated(green) versus varying 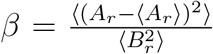.

Next, we simulate data under two conditions, where we keep the non-phase-locked statistics same for both the situations but vary the phase-locked activity statistics. We estimate the mean and variance of phase-locked and non-phase-locked amplitude via the CPM and averaging method. We infer the estimates from both the methods via t-test. As the non-phase-locked statistics are kept the same for both the conditions, we expect non-phase-locked estimates to fail the t-test, i.e., showing no statistical difference and phase-locked statistics to pass the t-test. **Tables 1 and 2** show the CPM and Averaging method results, respectively.

**Table 1.** Phase-locked and non-phase locked estimates and simulated values, using CPM.

| Case | Simulated Value | Estimated Value | T-test p-value | Case | Simulated Value | Estimated Value | T-test p-value |
| --- | --- | --- | --- | --- | --- | --- | --- |
| 1 $\langle A_r \rangle$ | 5.42 | 5.50 | | 1 $\langle B_r \rangle$ | 3.48 | 3.26 | |
| Var($A_r$) | 12.66 | 19.42 | **10<sup>-20</sup>** | Var($B_r$) | 5.9 | 0.6 | **0.10** |
| 2 $\langle A_r \rangle$ | 3.97 | 3.98 | | 2 $\langle B_r \rangle$ | 3.39 | 3.97 | |
| Var($A_r$) | 8.49 | 6.5 | | Var($B_r$) | 5.8 | 7.2 | |

**Table 2.** Phase-locked and non-phase locked estimates and simulated values, using Averaging method.

| Case | Simulated Value | Estimated Value | T-test p-value | Case | Simulated Value | Estimated Value | T-test p-value |
| --- | --- | --- | --- | --- | --- | --- | --- |
| Case1 $\langle pl\_power \rangle$ | 42.09 | 30.22 | | Case1 $\langle npl\_power \rangle$ | 18.04 | 29.44 | |
| Std($pl\_power$) | 48.62 | 49.60 | **10<sup>-14</sup>** | Std($npl\_power$) | 23.00 | 34.07 | **0.004** |
| Case2 $\langle pl\_power \rangle$ | 24.27 | 15.85 | | Case2 $\langle npl\_power \rangle$ | 17.36 | 25.26 | |
| Std($pl\_power$) | 32.98 | 32.67 | | Std($npl\_power$) | 22.08 | 30.35 | |

CPM method shows significant difference(p-value is 10^*−*20^) for phase-locked activity but not for non-phase-locked activity(p-value is 0.1). Whereas averaging method shows the significant difference for both phase-locked and non-phase-locked power with a p-value of 10^*−*14^ and 0.004, respectively. The averaging method gives inaccurate estimates as non-phase-locked power is affected by the statistics of phase-locked power, as shown in the previous section.

### 3.6 Application on Empirical Audio Oddball Data

We call the ratio 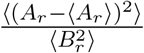 leaking parameter(*β*), if this ratio is less than 0.5, averaging method and CPM gives approximately similar estimates for NPL power but ratio greater than 0.5 CPM estimate should be used to calculate NPL power. In this section, we estimate the leaking parameter *β* from EEG audio oddball data. The EEG data was collected from an audio-oddball experiment; the audio stimulus presented to the subject is of two types, i.e., standard (frequent) and oddball(rare). In the experiment, the standard and oddball stimuli were the low frequency(261.6Hz) and high-frequency tones(523.3Hz). The experiment was conducted with 64 channel EEG with a sampling frequency of 1KHz. The data corresponding to each trial was 500ms in length(from the stimulus onset) and was re-referenced with the average. The band-pass filtering, artifact rejection, notch filtering at 50Hz(electrical transmission line frequency), and baseline correction were done to remove artifacts and increase the signal-to-noise ratio.

In **Figure 7** we calculate leaking parameter *β* for all the electrodes and frequencies (2 to 50Hz). The preceding section shows that *β <* 0.5, averaging and CPM estimates give approximately the same results. Since *β* varies over a long range, we represent *β* on log scale and use *β*= 0.5 as the critical value. Grey points in the **Figure 7** represents frequencies and electrodes for which CPM and averaging method gives same estimates, whereas aqua colored points represent frequencies and electrodes for which CPM estimate should be used, as averaging estimates are inaccurate.

**Figure 7:**
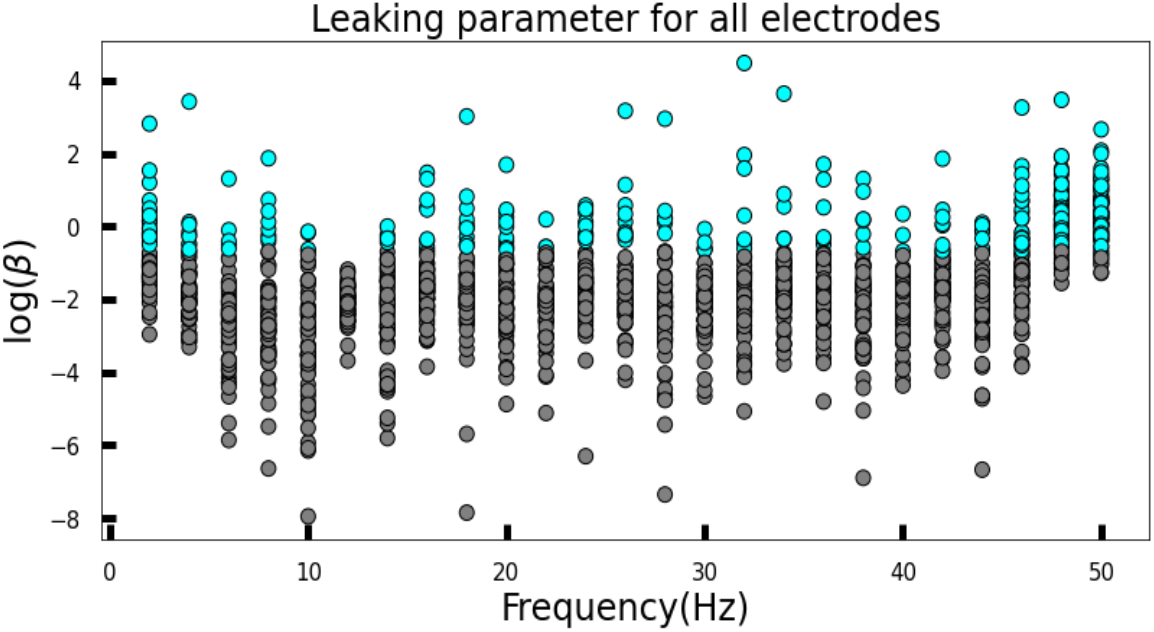
Leaking parameter(*β*) for audio oddball stimulus for all electrodes and frequencies. Aqua points are for *β>*0.5 and grey for *β<*0.5.

Next, we calculate NPL power at 2Hz for audio oddball EEG data using both CPM and the averaging method for each of 64 electrodes. We also calculate *β* i.e., the leaking parameter for each electrode. The log-log plot for estimated NPL power and *β* is shown in **Figure 8**. The averaging NPL power estimate is close to the CPM estimate for low *β* values, but CPM and averaging estimate differs significantly for electrodes with large *β* values. As *β* increases, the difference between averaging and CPM NPL estimate also increases. This shows that averaging method estimate deviates for electrodes with large *β* values and CPM estimates should be used for such cases.

**Figure 8:**
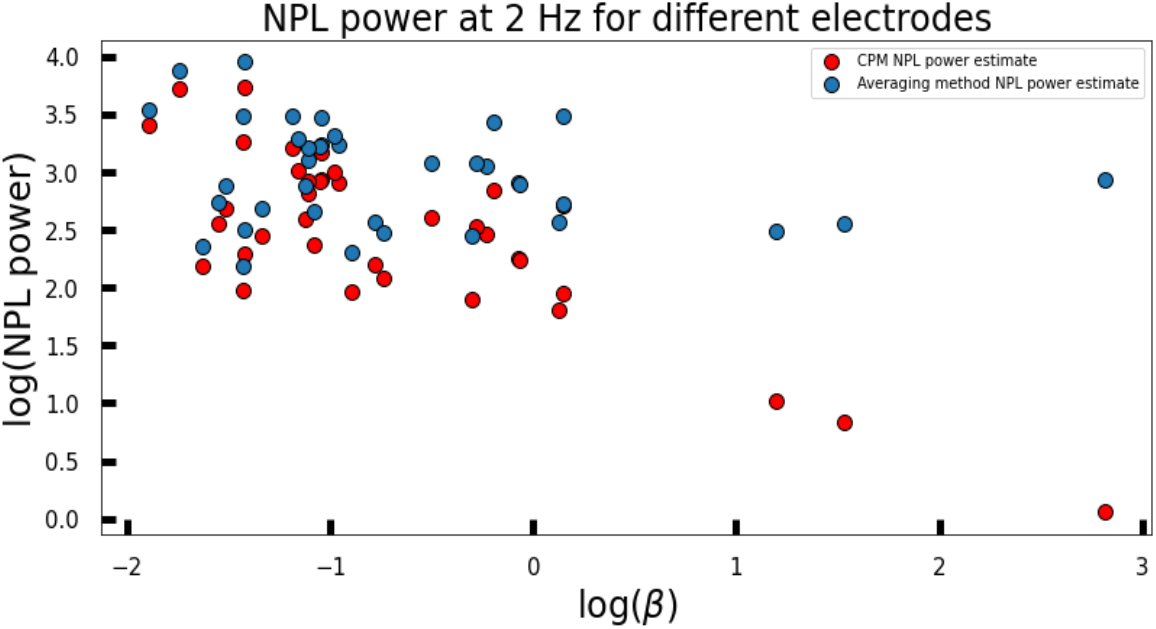
Comparison between averaging method and CPM estimate for NPL power. *log*(NPL power) estimated for each electrode at 48Hz versus *log*(*β*) calculated for each electrode.

### 3.7 Application on auditory steady state responses (ASSR)

Steady-state auditory stimulus is extensively studied, and its response is well established. Audio steady-state stimulus produces a response (called audio-steady-state response(ASSR)) which is close to the frequency at which stimulus is presented[Griskova-Bulanova et al., 2014]. It is used to test the ability of neuronal assemblies to synchronize at stimulus frequency. ASSR is predominantly reported to phase-locked response[Picton et al., 2003]. Here we test the established phase-locked nature of ASSR with our newly developed CPM.

A hundred trials of pure sin wave at frequency 500Hz, amplitude modulated at 40Hz with 100% modulation depth was presented to the subjects. The stimulus duration was of 1 second with 1-second inter-stimulus duration. The experiment was conducted with 63 channel EEG with a sampling frequency of 1KHz. The band-pass filtering at 2-48 Hz, artifact rejection, and baseline correction was done to remove artifacts and increase the signal-to-noise ratio. The signal was down-sampled to 250 Hz (to increase computation speed), and 2300 artifact-free trials of 1 sec ASSR were obtained.

As shown in the previous sections leaking parameter(*β*) tells how much PL power variation seeps into the estimate of NPL power in averaging method estimate. Here, the frequency of interest is 40Hz, at which we expect ASSR. We calculate *β* for the 63 electrodes at 40Hz to check if CPM estimates will differ from averaging method estimates. The results are shown in **Figure(9)**, panel [A] shows that *β* values are small for all electrodes(varying between −0.3 to 0.3), which is less than 0.5. Hence variance in *A*_*r*_ does not affect *B*_*r*_ estimates, as seen in the panel [B], where CPM and averaging method estimates of NPL power are approximately equal. **Figure (10)** shows that the NPL power remains at a pre-stimulus level, whereas PL power increases 10-80 times. These results align with the previously reported phase-locked nature of audio-steady state response.

**Figure 9:**
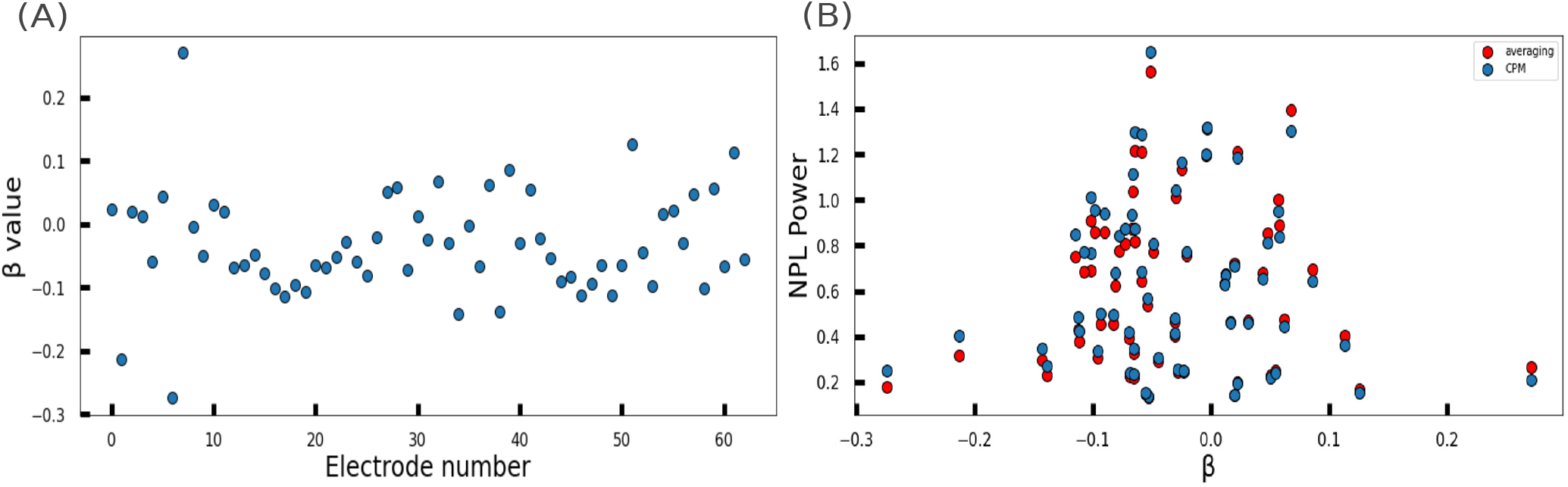
Leaking parameter estimate for each electrode to see if CPM estimates differ from averaging estimets. (A) Leaking parameter(*β*) for audio steady state stimulus for all electrodes at 40Hz frequency, (B) NPL power calculated for each electrode plotted against their *β* values, blue, red dots represent CPM, averaging method NPL power estimate respectively.

**Figure 10:**
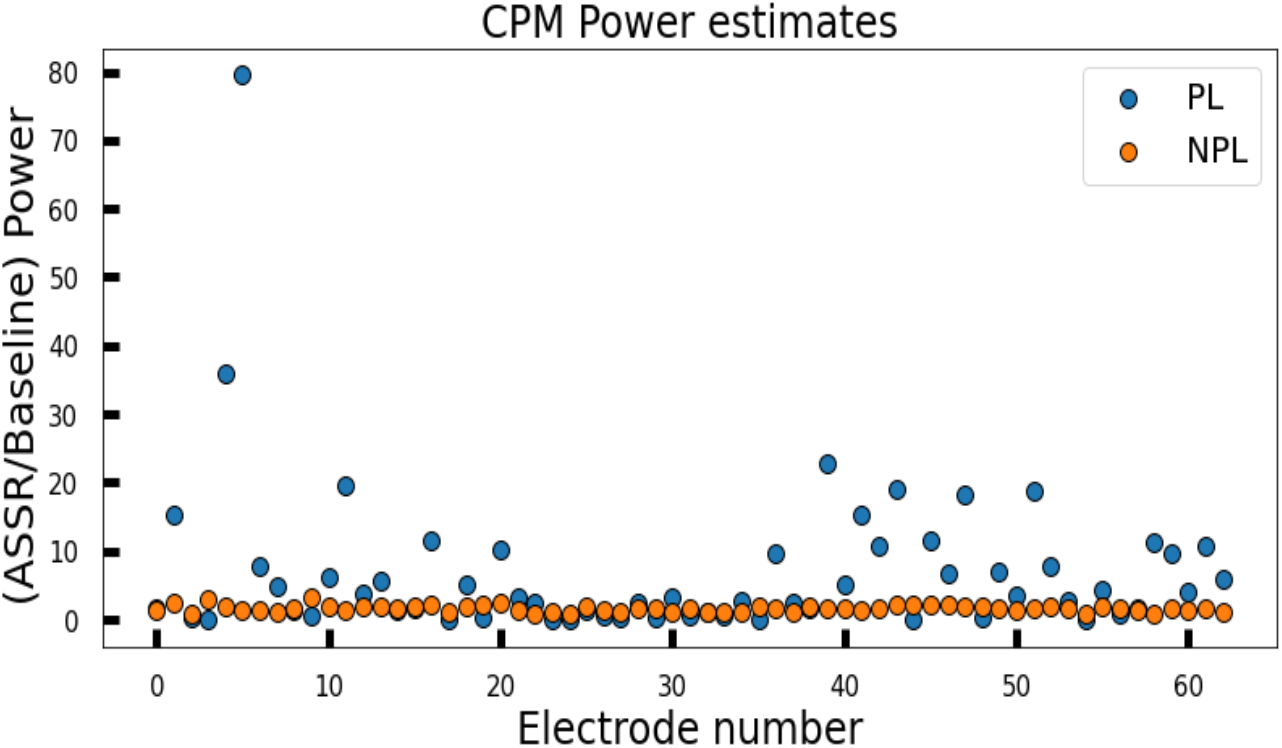
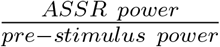 estimated via CPM for each electrode, blue dots gives PL power estimate relative to pre-stimulus power estimate and orange dots gives NPL power relative estimate.

## 4 Discussion

This article relaxed the assumption that the phase-locked activity is trial invariant(at a specific frequency). Studies like [McKewen et al., 2020], [Cohen and Donner, 2013] [van Noordt et al., 2022]raised the problem of variability in phase-locked activity corrupting NPL power estimate, but no method has been proposed to solve the issue. We consider both PL and NPL amplitude to vary with trials. We could not separate the PL and NPL activity at the single-trial level by proposing the CPM model. We proposed mean and variance estimators for PL and NPL amplitudes to make concise statistical inferences about the two activities. We observe that the CPM estimates approach averaging estimates for low PL variance. We applied the CPM on audio oddball EEG data and observed that *β* value was high enough for several frequencies and electrodes that lie outside the averaging method’s applicability. We recommend calculating the *β* ratio using CPM estimators to check the applicability of the averaging method. No variant of the averaging method concurrently considers both PL and NPL variation, making CPM a novel method. We applied CPM to estimate the PL and NPL power of ASSR in the audio-steady-state stimulus. We found *β* values to be low, thereby not affecting PL, NPL power estimates via the averaging method, and found CPM estimates to align with the previous results. We verified the phase-locked nature of ASSR using CPM. We recommend revisiting the established results, such as gamma oscillations being non-phase-locked and steady-state visual evoked responses being phase-locked [Tallon-Baudry and Bertrand, 1999] [Zhang et al., 2013]. Our CPM method correctly separates phase-locked activity from non-phase-locked activity at the group level.

An increase/decrease in NPL power estimates the degree of stimulus interaction with the background brain activity. A stimulus that elicits higher NPL power signifies higher interaction with the brain activity and thereby has a higher cognitive hierarchy than a stimulus that elicits lower NPL power. [Deiber et al., 2007] showed NPL activity was induced for detection tasks that require active working memory but not for the passive task. The study also shows that the induced power was larger for higher memory load. [Donner and Siegel, 2011] showed NPL activity are present in the state of arousal, affected by the level of cautiousness in decision making and sustained attention. An increase in NPL activity for experienced stimuli shows NPL activity corresponds to a complex interaction between stimuli and brain activity [Goffaux et al., 2004]. Increased gamma-band oscillations for recognized stimuli suggests that NPL activity subtend top-down influences of prior knowledge on bottom-up visual processing [Goffaux et al., 2004]. An increase in NPL activity is observed for the NOGO task that requires inhibitory response [van Noordt et al., 2022]. This study reiterates that NPL activity is enhanced when there is ongoing interaction between top-down and bottom-up processes.

Some studies have used the non-phase-locked activity to speculate the cognitive hierarchy in different stimuli processing. Though this hypothesis is not tested directly. A large number of reports shows enhanced NPL activity in higher-order cognition tasks [Jaušovec and Jaušovec, 2005]. This study shows that individuals with high emotional intelligence display more gamma-band NPL and smaller decrease in upper alpha band NPL than individuals with average emotional intelligence. Another proof that NPL is involved in higher order cognition is the study [Zhang and Han, 2021], which suggests NPL alpha oscillations are engaged in the spontaneous racial categorization of faces. All the mentioned studies suggest NPL activity can be used to probe the level of cognitive hierarchy of a stimulus. It will be interesting to directly test this hypothesis using a paradigm that parametrically controls cognition level.

Another point that needs further research is how stimuli interacts with background brain activity to give rise to trial-varying PL activity. We used CPM to show trial variation in phase-locked activity (verified by audio-oddball data). One can revisit the assumption that phase-locked activity is independent of ongoing background brain activity. The review paper by Nierhaus and colleagues [Nierhaus et al., 2009] discuss several studies showing that the evoked activity depends on the pre-stimulus background activity. Apart from trial-variability of evoked activity coming from the additive model, the phase-resetting model is proposed to be equally likely. No consensus could be drawn on the precise role of background activity or its relationship to evoked activity. In conclusion, single-trial separation of phase-locked and non-phase-locked power remains an open problem. Single-trial *A*_*r*_, *B*_*r*_ can be still be estimated approximately assuming *A*_*r*_, *B*_*r*_ to be Gaussian distributed and using the Bayesian approach to fit for *A*_*r*_, *B*_*r*_. A similiar approach is used by [Banerjee et al., 2012], [Bollimunta et al., 2007] to calculate single-trial lag and amplitude scaling factor for LFP and spike-field recordings. The single-trial calculation of evoked activity can shed light on the nature of the interaction between evoked and background brain activity.

## Acknowledgement

We thank Dr. Soibam Shyamchand Singh, Dr. Suman Saha, and Anagh Pathak for their feedback to improve the article’s readability. We also thank the Computing Facility of NBRC for infrastructure and resources.

## Appendix

### 4.1 Constant phase-*α* estimate

Taking trial average on both sides of equations (9) and (10) and assuming that *B*_*r*_ and *ϕ*_*r*_ are independent of each other gives

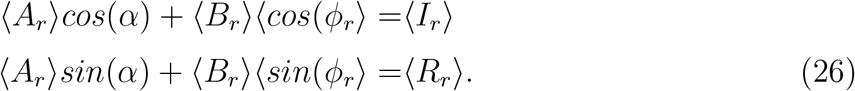

Since *ϕ*_*r*_ is expected to be uniformly distributed from −*π* to *π*, ⟨*cos*(*ϕ*_*r*_)⟩, ⟨*sin*(*ϕ*_*r*_)⟩ can be approximated to be zeros. This gives

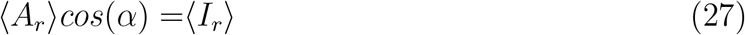

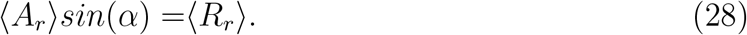

Dividing equations (27) by (28) gives

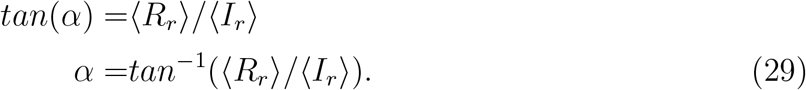

### 4.2 Trying other fourier coeffecients to solve for *A*_*r*_, *B*_*r*_, *ϕ*_*r*_

In section 3.2, we tried solving equation (3) for *A*_*r*_, *B*_*r*_, *ϕ*_*r*_, where we had two equations and three unknowns. Here we try creating more equations from equation(3). Hence, to solve for *A*_*r*_, *B*_*r*_, *ϕ*_*r*_ at single trial level, we require one more equation. Squaring *S*_*r*_(*t*), we get a non-zero dc component (i.e., Fourier coefficient at 0 frequency), and two equations corresponding to frequency at 2*ω*_0_. Taking fourier transform of 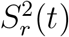 gives following equations

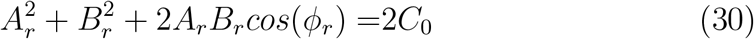

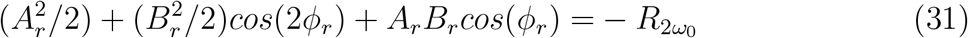

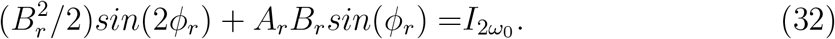

Where *C*_0_ is the Fourier coefficient at frequency 0, which is always real. 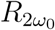 and 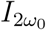 are the real and imaginary part of the Fourier coefficient at frequency 2*ω*_0_. Equations (30)-(32) and (9),(10) together gives us five equations. We can take any three equation to solve for three variables (*A*_*r*_, *B*_*r*_, *ϕ*_*r*_). We set to solve the above system of non-linear equations using a numerical method, for which we need Jacobian matrix of the above system of equations. Defining left hand side of equations as functions *f, g, h*, jacobian for 3×3 system is given by

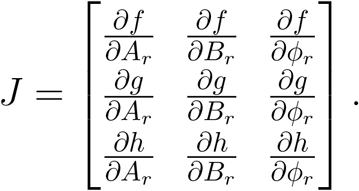

For the system of equations ((30)-(32)) Jacobian becomes

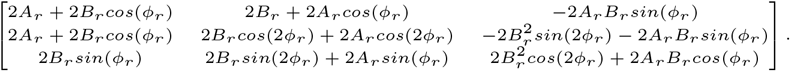

The determinant of jacobian is zero everywhere, which implies the inverse does not exist; therefore, we can not solve the system numerically. The determinant of jacobian is zero for any choice of three from five equations. Thus, the non-existence of a unique solution suggests that the phase-locked amplitude(*A*_*r*_) and non-phase-locked power(*B*_*r*_) can not be separated at the single-trial level.

### 4.3 The power operation

Here we define *Power*{.} as an operation that we used in equation 5. *Power*{.} is energy per unit time of the signal that it is operated on. It is given by sum of square of fourier coeffecient. Signal at right-hand side of 5 can be expanded as

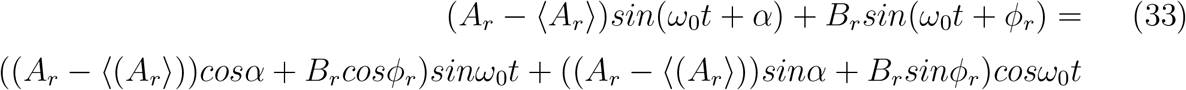

Operating power operation on both sides of 34, we get

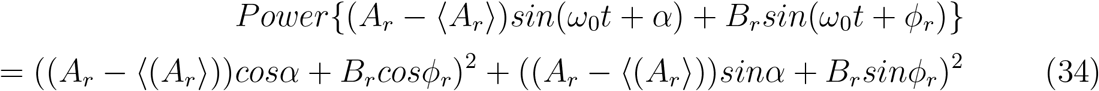

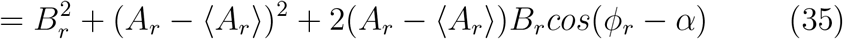

